# The dengue virus NS1 protein conveys pro-inflammatory signals by docking onto human high-density lipoproteins

**DOI:** 10.1101/2021.05.05.442454

**Authors:** Souheyla Benfrid, Kyu-Ho Park, Mariano Dellarole, James E. Voss, Carole Tamietti, Gérard Pehau-Arnaudet, Bertrand Raynal, Sébastien Brûlé, Patrick England, Xiaokang Zhang, Anastassia Mikhailova, Marie-Noëlle Ungeheuer, Stéphane Petres, Scott B. Biering, Eva Harris, Anavaj Sakunthabaï, Philippe Buchy, Veasna Duong, Philippe Dussart, Fasséli Coulibaly, François Bontems, Félix A. Rey, Marie Flamand

## Abstract

The nonstructural NS1 protein is a virulence factor secreted by dengue virus (DENV)-infected cells. NS1 is known to alter the complement system, activate immune cells and perturb endothelial barriers. Here we show that pro-inflammatory signals are triggered by a high affinity complex formed between NS1 and human high-density lipoproteins (HDL). Electron microscopy images of the NS1-HDL complexes show spherical HDL particles with rod-shaped NS1 protrusions on their surface. These complexes are readily detectable in the plasma of hospitalized dengue patients using anti-apolipoprotein A-I (ApoA-I) antibodies specific of the HDL moiety. The functional reprogramming of HDL particles by the NS1 protein as a means to exacerbate systemic inflammation during DENV infection provides a new paradigm linking the human lipoprotein network to dengue pathogenesis.

## INTRODUCTION

Dengue virus (DENV) infects nearly 400 million people annually, leading to more than 500,000 hospitalizations (*1, 2*). The mortality rate varies from less than 1% to 10% depending on the epidemic and medical care provided to patients (« Dengue and Severe Dengue » s. d. WHO, 2016, (*3*)). The dengue nonstructural protein 1 (NS1) is a viral effector circulating in the bloodstream of DENV-infected patients (reviewed in (*4–6*)). In DENV-infected cells, NS1 forms amphipathic dimers in the endoplasmic reticulum (ER) that insert into the luminal side of the membrane (*7–9*). The membrane-bound dimers play an essential role in orchestrating viral replication in specialized subcellular factories (*10*). A sub-fraction of NS1 dimers further associate by three to form barrel-shaped hexamers. During this process, NS1 hexamers detach from the ER membrane and are secreted as soluble nanoparticles loaded with lipids from the infected cell (*11, 12*). The secreted form of NS1 has previously been shown to bind to complement and coagulation factors, to activate immune and endothelial cells, to trigger the expression of pro-inflammatory cytokines, to alter the glycocalyx barrier and to promote endothelium permeability (*13–18*). The antibody response against NS1 has been shown to protect against several flavivirus infections (*14, 19–22*) but can also be harmful *via* a cross-reaction with platelets and endothelial cell surface antigens (*23–28*). These characteristics altogether favor the development of thrombocytopenia, vascular leakage and hemorrhage. Given the growing evidence of NS1 involvement in dengue pathogenesis, a better understanding of the molecular fate of NS1 in extracellular fluids by identifying its interacting partners is of utmost importance.

In the present study, we report that NS1 from dengue virus serotype 2 (DENV-2) binds high-density lipoproteins (HDL) and with a lower affinity low-density lipoproteins (LDL). HDL and LDL are lipoprotein complexes composed of large lipid cargos enwrapped by the apolipoproteins A-I and B, respectively, as well as a panel of functional proteins recently identified by proteomic approaches (*29, 30*). Lipoprotein particles that circulate in the blood have long been recognized for their regulatory functions in vascular homeostasis, inflammation and innate immune responses (*29, 31–34*). We explored the NS1-HDL association by biophysical methods and visualized the complex by electron microscopy. Our observations revealed a stable anchoring of NS1 amphipathic dimers to the HDL surface. We observed that the NS1-HDL complex could trigger the production of pro-inflammatory cytokines in exposed human primary macrophages. In addition, we consistently detected elevated levels of NS1-HDL complexes in the blood of DENV-infected patients on the day of admission to hospital using an anti-apolipoprotein A-I (ApoA-I) detection assay. We further determined that NS1 complexes acquired an apolipoprotein E (ApoE)-positive phenotype during the clinical phase, a component mostly found on very-low-density lipoproteins or chylomicrons and only transiently associated to HDL or LDL. This pointed to a complex and dynamic interaction of NS1 with the host lipoprotein metabolic cycle.

## RESULTS

### The DENV-2 NS1 hexamer binds high- and low-density lipoprotein particles

In this study, we first sought to identify NS1 protein partners encountered during its circulation in human blood. For this purpose, we carried out a pull-down assay using a purified preparation of recombinant strep-tagged DENV2 NS1 spiked in plasma obtained from healthy donors to pull down potential ligands. We thus re-affinity purified NS1 from the plasma and analyzed the resulting products by size exclusion chromatography (SEC) (Fig. 1A). Compared to NS1 alone, the pull-down SEC profile showed an additional peak and a large shoulder at smaller elution volumes, corresponding to apparent molecular weights of 840 and 380 kDa, respectively (Fig. 1A). The protein contents of the two high molecular weight complexes were analyzed by SDS-PAGE and the identities of the predominant protein bands determined by N-terminal sequencing and mass spectrometry as ApoA-I and ApoB. These proteins correspond to the main scaffold proteins of HDL and LDL, respectively (Fig. 1A).

**Figure 1:**
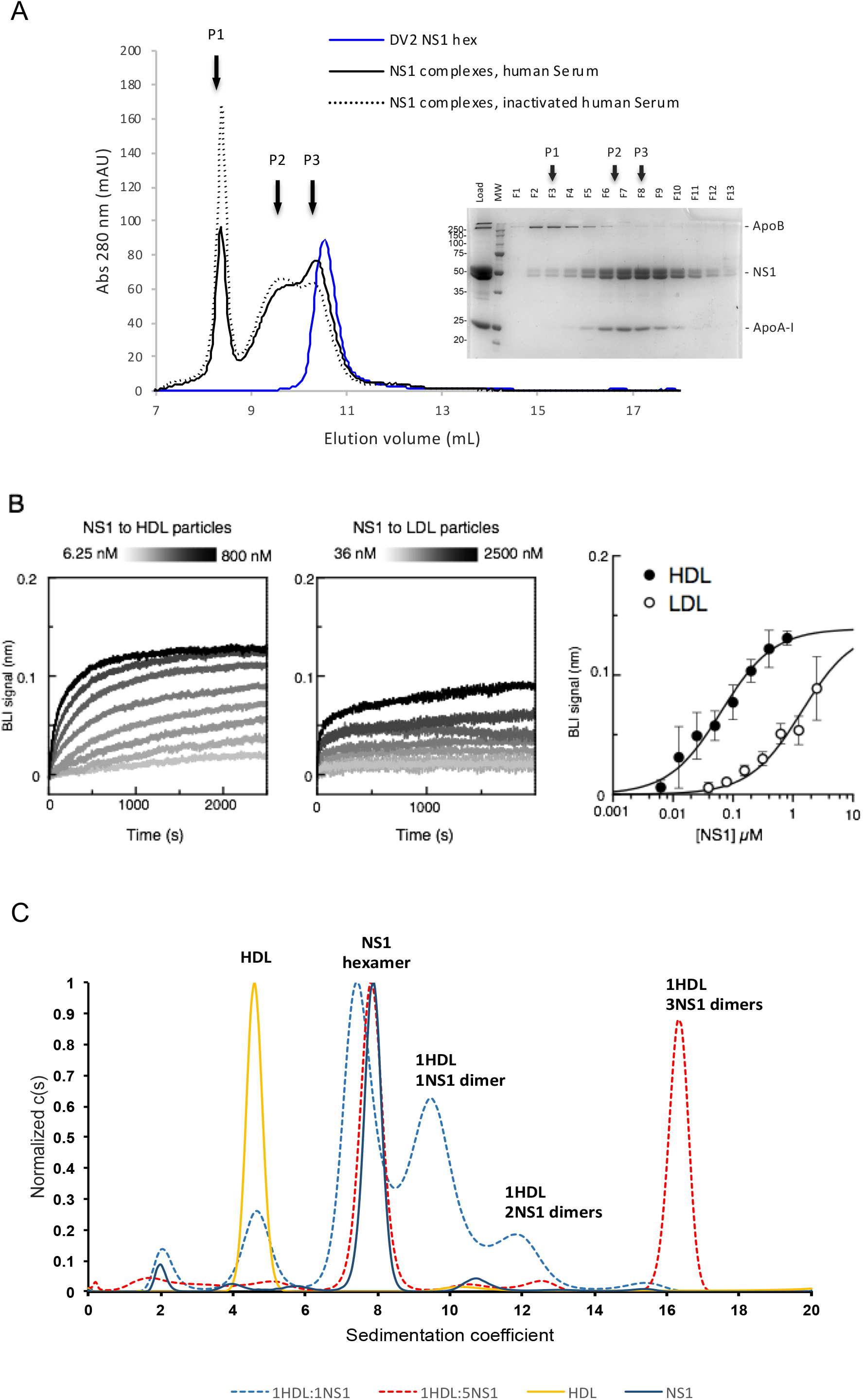
DENV NS1 binds preferentially to human high-density lipoproteins. (A) Size exclusion chromatography profile of NS1 pull-down experiments showing a clear shift after incubation in complete or heat-inactivated human serum (solid and dotted black lines, respectively) from the same healthy donor compared to the NS1 protein alone (blue line). NS1 protein partners were identified by SDS-PAGE and N-terminal sequencing as the Apolipoprotein B scaffold of the low-density lipoproteins (LDL) in the first SEC elution peak, and the ApoA-I protein scaffold of the high-density lipoproteins (HDL) in the second elution peak. (B) Biolayer interferometry (BLI) profiles corresponding to the binding of serial-diluted concentrations of NS1 respectively to human HDL (left panel) and human LDL (central panel) particles. The concentration-dependence of the steady-state signal corresponding to the binding of NS1 to HDL (black dots) and LDL (white dots) is shown on the right hand side panel. (C) Typical sedimentation coefficient distribution of NS1 or human HDL alone or pre-incubated together (mix of NS1 and HDL at a 1:1 or 5:1 mass ratio) monitored using an interferometric detector. Peaks were integrated for all the detectors (interference and absorbance at 280nm). The calculated stoichiometries are indicated for each peak. Solutions were equilibrated at 20°C for 2h before sedimentation velocity analysis.

This observation prompted us to assess the affinity of DENV2 NS1 for HDL and LDL. We immobilized human HDL and LDL particles on bio-layer interferomety (BLI) sensors coated with specific polyclonal antibodies against ApoA-I or ApoB, respectively. Figure 1B displays the binding curves for increasing NS1 concentrations in contact with both types of lipoprotein particles, and the values reached at steady state. The curves can be fitted using a single state binding model leading to a relative binding constant (Kd) of 63 nM ± 0.2 nM for HDL and 1.4 μM ± 0.1 μM for LDL (Fig. 1B), indicating preferential binding of NS1 to HDL compared to LDL.

In order to characterize the architecture of the complex, we used analytical ultracentrifugation to study the behavior of NS1, HDL and a mix of NS1 and HDL at either a 1:1 or 5:1 mass ratio, which represents approximately a 0.5:1 or 2.3:1 NS1 hexamer to HDL molar ratio (Fig. 1C). Purified NS1 sedimented as a main species with a sedimentation coefficient of 7.9 S compatible with a globular hexamer. HDL particles exhibited a much lower value of 4.20 S in keeping with the larger lipid to protein ratio (*35*). In the sample containing an excess of NS1 mixed with HDL, all the HDL was engaged in an interaction with NS1 forming a unique species with a sedimentation coefficient of 16.8 S that segregated distinctly from the other species (Fig. 1C). Residual unbound NS1 hexamer could still be observed, as expected due to the NS1 excess (Fig. 1C). By combining two detectors and taking into account the theoretical composition of the HDL particles, we estimated that one NS1 hexamer was bound to each HDL particle when present in excess. Interestingly, in the 1:1 ratio sample, we detected intermediate peaks at 9.4 and 11.7S that corresponded to one or two NS1 dimeric subunits bound to HDL, respectively.

### Binding of the NS1 hexamer to spherical HDL leads to a surface rearrangement of NS1 into dimeric rod-shaped protrusions

Based on the above results, we used a 2:1 molar ratio to prepare NS1-HDL complexes and examined the resulting products by negative stain electron microscopy (EM). As previously shown (*36*), human HDL particles appear as smooth spheres ~10 nm in diameter with an electron-dense central region (Fig. 2A). The purified NS1-HDL complexes, in contrast, presented a granular surface with prominent structures on their outer surface (Fig. 2B). 2D class averages of NS1-HDL complexes revealed that the HDL particles were crowned with high-density features that match very well the contour of NS1 dimers in side view with two discrete nodules that likely correspond to the NS1 protomers (Fig. 2B, 2C) (*7–9*). About 60% of the complexes analyzed in our study presented three NS1 dimers on the HDL surface, suggesting that NS1 hexamers dissociate into dimers upon binding to HDL particles (Fig. 2B, 2C). Around 25% and 10% of the particles presented 2 or 4 apparent NS1 dimers at their surface, respectively. These observations corroborate the ultracentrifugation data showing different ratios of NS1 dimers per HDL particle depending on the initial NS1:HDL ratio (Fig. 1C) and suggest a dynamic binding mode between NS1 and HDL particles.

**Figure 2:**
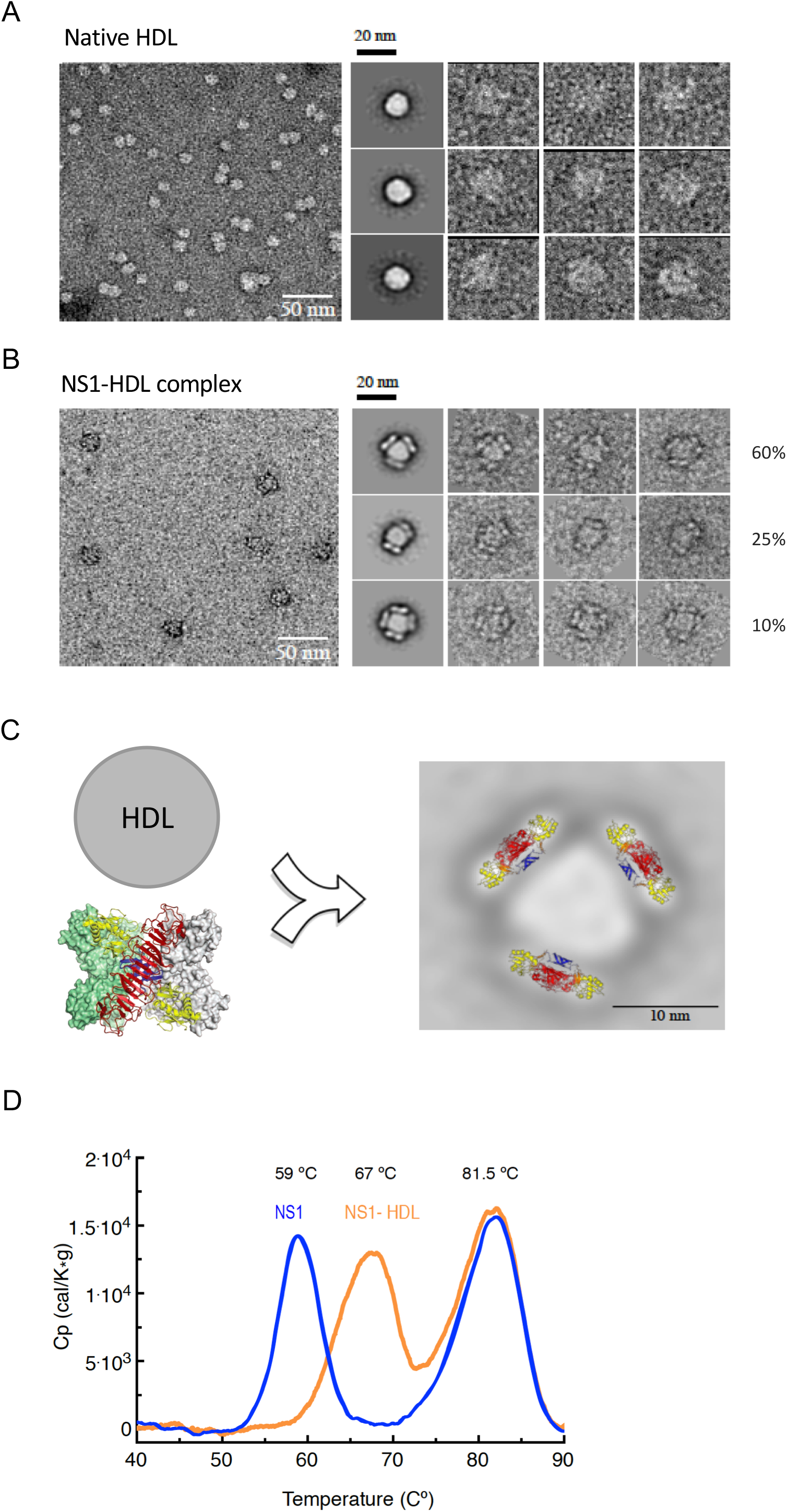
Analysis of NS1-HDL complexes by electron microscopy reveals the presence of NS1 dimers at the surface of HDL particles. (A, B) Electron microscopy observations from left to right: a representative image, followed by the three most representative classes of purified HDL particles (A) and NS1-HDL complexes (B). (C) Fitting of the NS1 3D structure of the dimeric form into the most abundant class of NS1-HDL complexes, suggesting a collapse of the NS1 hexamer into hydrophobic dimeric blocks that float and anchor into the HDL lipid phase. (D) Differential scanning calorimetry (DSC) of NS1 alone (blue line) or of an NS1-HDL mix at a 2:1 molar ratio (orange). Of note, the HDL particles alone did not generate any signal in the temperature range tested.

The presence of NS1 dimeric subunits associated to HDL was also evidenced by DSC (Fig. 2D). Thermal scanning of both hexameric NS1 and the NS1:HDL mix (at a 2:1 molar ratio) showed two transitions phases while the HDL particles alone did not contribute to any signal in the scanned temperature range (Fig. 2D) (*37*). The second transition at a Tm of 81°C was identical for NS1 alone and in complex with HDL. We have previously reported that the NS1 dimer of Japanese encephalitis virus requires temperatures higher than 80°C to dissociate into monomers (*38*). We therefore attributed this second transition to the dissocation and full denaturation of NS1 dimeric subunits (Fig. 2D). Accordingly, the first transition peak corresponds to the dissociation of NS1 hexamers into dimers for NS1 alone (Tm of 59°C) and to the release of NS1 dimers from the HDL particle for the NS1-HDL mix (Tm of 67°C) (Fig. 2D). The difference in Tm values observed for the first transition peak in both samples provides additional evidence that once the NS1-HDL complex formed, the NS1 dimer-dimer interface initially present in the hexamer is converted into a more stable interface formed between the NS1 dimers and the HDL surface.

### The NS1-HDL complex triggers pro-inflammatory signals in human primary macrophages

NS1 is known to trigger the production of pro-inflammatory cytokines in macrophages (*15*). As HDL are potent modulators of inflammation (*31, 32*), we characterized the cytokine and chemokine production pattern in human macrophages exposed to NS1 alone, HDL alone or to the NS1-HDL complex (Fig. 3B). Primary macrophages were differentiated from isolated monocytes of various donors and stimulated with the different combinations of effectors. When exposed to NS1 or HDL alone, we observed no significant difference in the cytokine levels produced by macrophages compared to the negative control, whereas the bacterial lipopolysaccharide (LPS) consistently induced high levels of cytokine secretion (Fig. 3B). These observations rule out any cytotoxic effect from putative contaminants in the NS1 and HDL samples. In contrast, the NS1-HDL mix induced a significant increase in TNF-α, IL-6 and IL-10 secretion, and to a lesser extent Il-1β, compared to NS1 alone. The differences were even higher when compared with HDL alone, suggesting that NS1 converts HDL into pro-inflammatory signaling particles (Fig. 3B).

**Figure 3:**
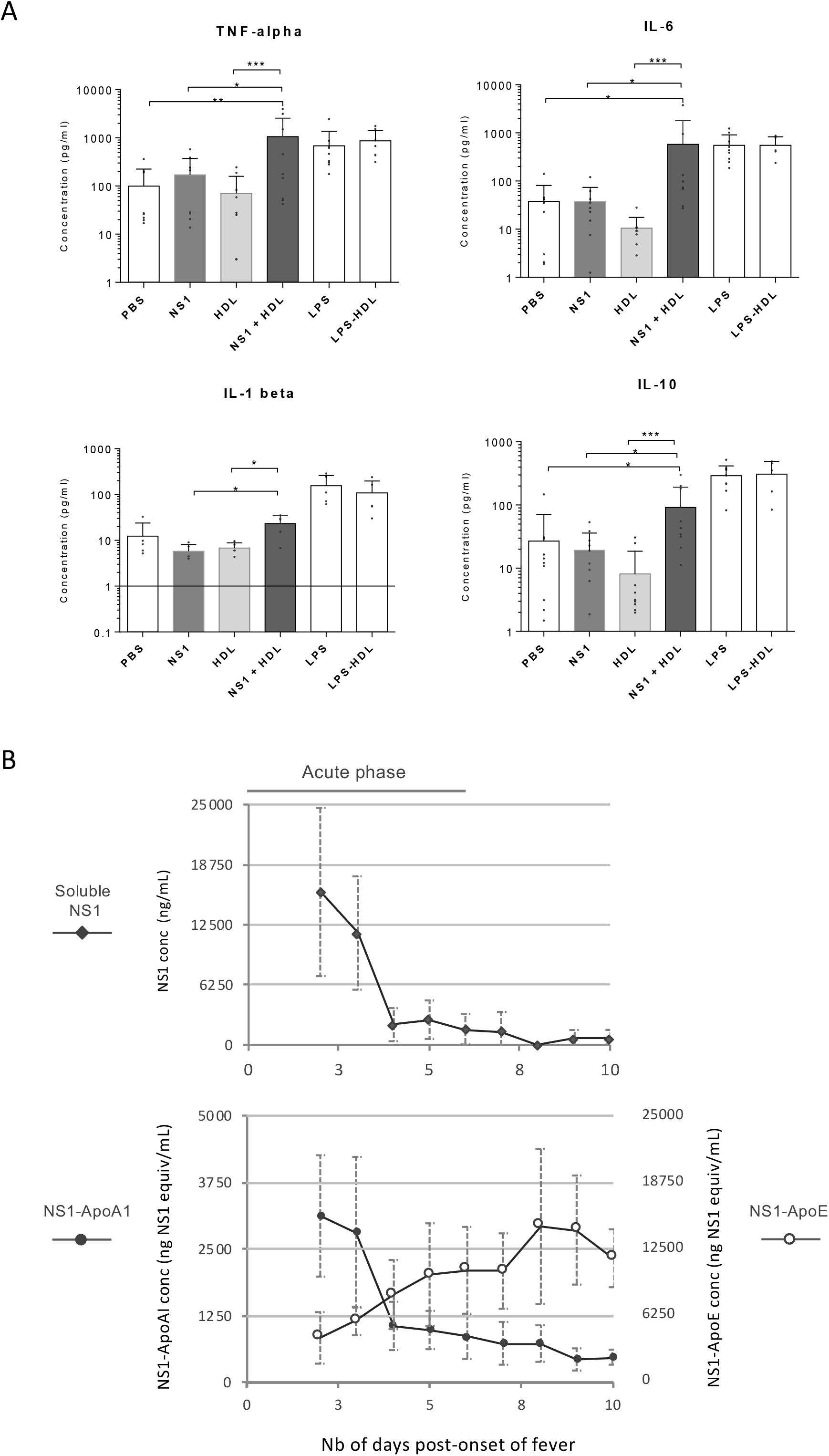
The NS1-HDL lipoprotein complex is biologically active in human primary macrophages and is detectable over the clinical phase in DENV-infected patients. (A) The NS1-HDL complex activates the inflammatory cytokine production from human primary macrophages. After 24h incubation with respective effectors, supernatants of cell culture were recovered and Luminex assays were performed to quantify the amount of TNF-alpha, IL-6 and IL-10 released by cells in the extracellular medium. Data represent mean +/− SEM of three independent experiments obtained using separate cells from four donors. LPS stimulation was used as a positive control in the presence or absence of HDL and provided values consistent between experiments. 2-Way analysis of variance (ANOVA) multiple comparisons were used to assess the statistical significance of differences observed between mean cytokine levels in culture supernatants between the different conditions. (B) Patients (n=51) hospitalized in the Kampong Cham Referral Hospital, Cambodia, presented either dengue with warning signs or severe dengue. A blood sample was recovered for each patient on the day of admission at hospital and during a follow-up visit that occurred before discharge from hospital (on average 4 days apart). Both samples were tested for the presence of the NS1 protein (upper panel), the NS1-ApoA-I complex or the NS1-ApoE complex (lower panel, dark and white dots, respectively) using three different ELISA formats as described in Supplementary Fig. 1. NS1-ApoA-I complexes are representative of the NS1-HDL particles in plasma samples. NS1-ApoE may include different types of NS1-lipoprotein complexes as ApoE is an exchangeable lipropotein that binds to various types of lipoprotein particles at different stages of their metabolic cycle.

### NS1-HDL complexes are detected in hospitalized patients

Knowing that the concentration of the above-mentioned cytokines and chemokines is dramatically increased in patients with severe dengue (*39–42*), we assessed the presence of NS1-HDL complexes in DENV-infected patients. To this end, we developed an additional ELISA format to broadly detect and quantify NS1-lipoprotein complexes in human plasma. In addition to our original NS1 detection assay (*43, 44*) and to the NS1-HDL semi-quantitative ELISA (Supplementary Fig. 1A,1B), we used an anti-ApoE PAb for the secondary detection of NS1 complexes potentially formed with the various types of lipoprotein particles (VLDL, IDL, chylomicrons in addition to HDL and LDL, as reviewed in (*45, 46*)) (Supplementary Fig. 1C). Using the three different ELISA formats, we tested blood samples from 51 dengue patients starting on the day of admission to hospital until their discharge. This represented on average a time lag of 4.3 days between the first and last samples. The vast majority of patients (around 80%) showed significantly elevated NS1 and NS1-HDL in blood on the day of admission compared to the last sample (Fig. 3C). The highest levels were observed between day 2 and day 4 post-onset of fever (Fig. 3C). By contrast, concentrations of NS1-ApoE-positive lipoprotein particles increased over time and were highest when tested on the last blood specimen recovered at the time of discharge from hospital (Fig. 3C). At this stage, we do not know whether the NS1-HDL complex recruits the exchangeable ApoE as a maturation or recycling process or whether, once the HDL pool becomes exhausted, the NS1 protein still expressed binds to other types of lipoprotein particles. In any case, our data clearly show a transition in the nature of the complexes formed between NS1 and the host lipoproteins during the course of infection.

## DISCUSSION

Dengue virus nonstructural protein 1 is a viral virulence factor that contributes to the development of severe dengue, characterized by cytokine storm, thrombocytopenia, vascular leakage and hemorrhage (*4–7*). NS1 circulates in the blood of infected patients at nanogram to microgram per ml levels (*47–49*). We previously described that the secreted hexameric form of NS1 shuttles a lipid cargo analogous to those carried by plasma lipoproteins, suggesting a possible interference with the lipoprotein metabolic cycle (*12*). Herein, we provide evidence of a direct interaction between NS1 and human HDL characterized by the ApoA-I scaffold protein, and with LDL identified by the presence of ApoB. The complexes formed between NS1 and HDL accumulate during the acute phase of the disease and by the end of the hospitalization period, NS1-lipoprotein complexes acquire an ApoE-positive phenotype that may be part of the anti-viral response engaged by the host.

Once bound to an HDL particle, the NS1 hexamer appears to fuse and collapse into NS1 dimeric building blocks that sit on the surface of the lipoprotein particle. It is not clear at this stage to which extent protein-lipid or protein-protein interactions prevail but both are likely to be important. Indeed, NS1 dimers are known to have the ability to bind lipid membranes and liposomes (7-9) and all the candidate protein ligands of NS1 published so far belong to the HDL proteome (*50*). These proteins include complement factors C4, C1s, hnRNP C1/C2, factor H, prothrombin, as well as inhibitory factors of complement clusterin, C5-9 and SC5b-9 complexes (*51–55*). NS1 does not necessarily interact with all but it is reasonable to anticipate that some will favor the association of NS1 with HDL or its stabilization. Proteomic studies, direct protein-protein interaction assays, the use of synthetic HDL with defined compositions or alternatively mimetic peptides will be instrumental in delineating the role of the different NS1 partners in the formation of a biologically relevant NS1-HDL complex.

NS1 was previously shown to activate the immune system and impair vascular homeostasis. NS1 can trigger the production of inflammatory cytokines and chemokines in cell culture and in immunodeficient mice (*14, 16, 56, 57*). Our findings demonstrate that the formation of NS1-HDL particles is a prerequisite for this effector function. The NS1-HDL complex assembled *in vitro* triggers pro-inflammatory signals in primary human macrophages while NS1 or HDL alone have no comparable effect. HDL particles are known to have an anti-inflammatory regulatory function and contribute to the maintenance of vascular integrity under physiological conditions (*29, 31–33*). However, it has also been reported that the recruitment of specific proteins by HDL, such as serum amyloid A (SAA), confers a pro-inflammatory status to these particles during an acute phase response (*58–62*). Our working hypothesis is that NS1 exerts a similar control on HDL during dengue virus infections. This process could involve the HDL scavenger receptor B1 that has recently been identified as a cell surface receptor for dengue virus NS1 (Alcalà et al., BioRixV, under review) allowing its internalization in many mammalian cell types including macrophages, endothelial cells, keratinocytes and hepatocytes. An NS1-HDL contribution to the cytokine storm would have a direct impact on the development of severe dengue as increased levels of TNFα, IL6 and IL-10 correlate consistently with disease severity and in certain instances endothelium permeability (*63–69*).

Several studies have described altered levels of HDL, LDL or VLDL in severe dengue (*70–75*). It is conceivable that NS1 broadly impacts the lipid and lipoprotein pathways by reprogramming the signaling patterns of the different lipoprotein particles or modulating the rate of their metabolic turnover. In this respect, we observed a notable change in the composition of the NS1-bound lipoprotein particles with the acquisition of an ApoE-positive signature over the course of the disease. ApoE is an exchangeable lipoprotein that can associate with most lipoprotein particles during the lipid metabolic cycle (*45, 46*). ApoE has also been recognized for its anti-inflammatory, anti-oxidative, anti-thrombotic and endothelial repair related properties (*76, 77*). The changes we observe emphasize the dynamic nature of the interaction of NS1 with host lipoproteins. Further studies are now needed to investigate whether the formation of NS1-ApoE-positive lipoprotein complexes are part of a recovery mechanism from the host or whether NS1 continues its intoxication program during the convalescent phase. A number of studies have described a persistence of asthenia for weeks, extending well beyond the end of the acute clinical phase (*78–81*).

In conclusion, we find that the secreted form of DENV protein NS1 hijacks HDL from human serum, and possibly other lipoprotein particles such as LDL, to form a pro-inflammatory NS1-HDL lipoprotein complex. Our data further suggest the occurrence of a structural transition of NS1 on the surface of HDL that results in the collapse of NS1 hexamers and the anchoring of amphipathic dimeric subunits in the HDL lipid phase. Finally, we demonstrate the presence of NS1-HDL complexes in the sera of dengue patients during the acute phase of the disease. The capture of HDL by NS1 may represent a way for the virus to exacerbate systemic inflammation. This will affect vascular permeability and facilitate virus propagation in the infected organism. Unraveling the molecular mechanisms governing the assembly of the NS1-HDL complex, its metabolic fate and pathogenic functions will be critical in defining preventive measures against the active form of DENV NS1.

## MATERIALS AND METHODS

### DENV-2 NS1 protein expression, purification and serum pull-down experiments

The DENV-2 recombinant NS1 protein was expressed in drosophila S2 cells and purified from the extracellular medium as detailed in the previously published supplementary methods (*12*). Purified DENV-2 recombinant NS1 protein was incubated for 1h at 37°C in serum or plasma recovered from a healthy donor (provided by the ICAReB facility, Institut Pasteur) at a final concentration of 400 μg NS1/mL plasma. The mix was then purified on a Strep-tactin column (Iba), washed twice with PBS Mg^2+^/Ca^2+^ (Gibco) followed by 14 column volumes of PBS 0.3 M NaCl and another 5 column volumes of PBS Mg^2+^/Ca^2+^. Elution was performed using 2.5 mM D-desthobiotine (Iba) in PBS Mg^2+^/Ca^2+^.

Purified samples of recombinant NS1, human HDL (Millipore), human LDLs (Millipore) or an *in vitro* reconstituted NS1-HDL mix were analyzed by size exclusion chromatography on a Superdex 200 10/300 column (GE healthcare). Protein standards from Bio-Rad were used to interprete elution profiles. The protein samples from the major peaks were further denatured in 5x Laemmli sample buffer containing β-mercaptoethanol, boiled for 5 min at 95 °C and separated by discontinuous sodium dodecyl sulfate (SDS) 4-15% polyacrylamide gel electrophoresis (SDS-PAGE precast gels, Bio-Rad). The SDS-PAGE gels were stained in Coomassie Blue solution (Bio-Rad).

### Biolayer Interferometry

DENV2 NS1 binding to HDL and LDL particles was monitoreded by Biolayer Interferometry (BLI), using an Octet Red384 instrument (ForteBio). Streptavidin-coated biosensors (SA, ForteBio) were loaded with biotinylated anti-ApoA-I or anti Apo-B antibodies (Abcam), followed by HDL or LDL, respectively. Subsequently the biosensors were incubated in wells containing serial dilutions of NS1 protein (concentrations ranging from 6.25 to 800nM for HDL, and from 36 to 2500nM for LDL) and the BLI association signals were recorded in real-time until they reached a plateau. Finally, the biosensors were incubated in wells containing buffer to monitor the dissociation of the complexes formed, before being regenerated for further use in replicate experiments. The regeneration protocol, comprising three subsequent 20 seconds washes in 10 mM Gly-HCl pH2, could be applied up to eight times for up to two days without losing any loading capacity of the immobilized biotinylated antibodies. The specific NS1 binding curves were obtained by subtracting the non-specific signals measured on unloaded biosensors used as control references. The steady-state signals were determined at the end of the association step and fitted using the following equation: Req= Rmax*C/Kd+C where Req is the steady-state BLI response, C the NS1 concentration, and Rmax the response at infinite concentration. All measurements were performed at least three times to determine experimental errors. All experiments were performed at 20°C in PBS Mg^2+^/Ca^2+^ (Gibco) supplemented with 0.1% milk to minimize nonspecific binding, using 96-well half-area plates (Greiner Bio6One) filled with 150 μl per well, and a shaking speed of 1000 rpm. Data was processed using the Scrubber (v2.0 BioLogic), BIAevaluation 4.0 (Biacore) and Profit (Quantumsoft) softwares.

### Analytical ultracentrifugation

NS1, HDL, and NS1-HDL mix at different molar ratios were centrifuged at 32,000 rpm for the complexes in a XL-I and a new Optima AUC (Beckman Coulter) analytical ultracentrifuge, at 20 °C in a four-hole AN 50–Ti rotor equipped with 3-mm and 12-mm double-sector aluminium epoxy centrepieces.

Detection of the biomolecule concentration as a function of radial position and time was performed by absorbance measurements at 250 nm and 280 nm, and by interference detection. Ultracentrifugation experiments were performed in PBS+/+ (Gibco). Sedimentation velocity data analysis was performed by continuous size distribution analysis c(s) using the Sedfit 15.0 software (*82*). All the c(s) distributions were calculated with a fitted fractional ratio f/f0 and a maximum entropy regularization procedure with a confidence level of 0.95. Buffer viscosity and density as well as the extinction coefficient of NS1 were calculated using the sednterp software (http://sednterp.unh.edu/). Partial specific volume were measured for NS1 (0.82 ml.g^-1^, as defined on an iodixanol density gradient) and confirmed by multidetection AUC experiment for HDL (0.94 ml.g^-1^). Extinction coefficients were extrapolated at 250 nm using the Utrascan II software. Deconvolution of the multi detectors signal into stoichiometric ratio was performed by integrating all the peaks on each detector to determine the contribution of each partner present (NS1, HDL or both).

### Differential scanning calorimetry (DSC)

Thermal unfolding of NS1 and of the NS1-HDL complex were followed using a VP-Capillary DSC instrument (Malvern MicroCal) in PBS buffer. The concentration of the NS1 hexamer was 0.2 mg/ml and was used at a 2:1 molar ratio to form the NS1-HDL complex. At least two DSC scans were recorded for each sample. Human HDL (Merck Millipore) was incubated with NS1 for 1 hour at 37°C prior to the DSC experiments. Scan rate was 100°C/hr with a filtering period of 2. Thermograms were analysed with the Origin software provided by the manufacturer.

### Electron microscopy and image analysis

Solutions of NS1 and NS1-HDL were spoted on glow discharged carbon grids, contrasted with 2% uranyl acetate and imaged with a Tecnai F20 microscope (Thermo Fisher, USA) in low dose conditions. Automated acquisitions were performed using EPU software (Thermo Fisher, USA) and images were acquired usind a Falcon II (Thermo Fisher, USA) direct detector.

HDL and NS1-HDL images were CTF-corrected (phase flip) and sorted using the XMIP software (*83*). Corrected images were imported in Relion (*84*). The recommended strategy for particle picking was applied as follow: a manual selection of particles compatible with the HDL or NS1-HDL size was performed on a small number (about fifteen) of images. A 2D classification (40 classes) was performed and five representative well-defined classes were selected as template for the automatic picking, leading to about 30 000 particles. A 2D classification (200 classes) was then performed. Classes obviously corresponding to artefacts were suppressed and a final run of 2D classification (200 classes) was carried out.

### Macrophages immune activation assay

Monocytes were isolated from buffy coats and were differentiated into macrophages by using human AB serum in macrophage medium, as previously described (Allouch et al., PNAS 2013). Briefly, PBMCs were isolated from whole blood using a Ficol gradient centrifugation (Eurobio). CD14+ cells are purified by magnetic bead separation of PBMCs using CD14+ human positive selection kit (StemCell) and plated 1M cells per ml on teflon plates (Sarstedt) with 7 ml per plate in the following medium: RPMI-1640 (Gibco), 2 mM L-glutamine (LifeTechnologies), 1% penicillin-streptomycin of concentration: 10,000 units penicillin and 10 mg streptomycin/ml (LifeTechnologies), 10 mM Na Pyruvate (LifeTechnologies), 10 mM HEPES (LifeTechnologies), 1% MEM vitamins (LifeTechnologies), 1% NEAA (LifeTechnologies), 50 uM beta-mercaptoethanol (LifeTechnologies), 15% human serum (ICAReB facility, Institut Pasteur). Monocytes were cultured in differentiating media for 6-8 days after which the macrophages were scraped off Teflon plates and counted. After spinning, they were resuspended at 1 million/ml in the same media but with 5% FBS instead of human serum.

Macrophages were plated at 0.5 million cells per mL in P24 plates (Corning). After 2 hours sedimentation and adhesion of the cells, macrophages were incubated with PBS (as negative control), LPS (100 ng/mL) as positive control, HDL (2.5 μg/mL), NS1 (10 μg/mL) or a NS1-HDL mix at same respective quantities (2:1 molar ratio) for 24h before collection of the supernatants. Inflammatory mediators were detected in cell supernatants using a hMagnetic Luminex Assay 5 Plex, R&D Systems, Bio-Techne Ltd run on the BioPlex 200 System xMAP (BioRad Laboratories Inc.) as per manufacturer’s specifications. The antibody bead kit was designed to quantify IL-1β, IL-6, IL-10, TNF-alpha. Standards were run with each plate at every assay to titrate the level of cytokines present. Statistical analyses were performed in Prism 6.0 (GraphPad Software Inc.) Data are shown as individual points and means ± SD. Significant testing was performed using 2-way ANOVA.

### DENV-infected patient sera specimen

Patients presenting acute dengue-like symptoms – between June and October of 2011 and 2012 – were enrolled at the Kampong Cham Referral Hospital. Inclusion criteria, following the WHO 1997 classification scheme, were children between 2 and 15 years old who had fever or history of fever at presentation and onset of at least two of the following symptoms within the previous 72 hours: headache, retro-orbital pain, muscle pain, joint pain, rash, or any bleeding signs. We performed a prospective, monocentric, cross-sectional study of hospitalized children with severe and non-severe dengue. The study was approved by the Cambodian National Ethics Committee for Human Research (approval #087NECHR/2011). All patient inclusion and blood sampling occurred after obtaining written informed consent from the patient’s parents or guardians. The first visit was conducted at hospital admission. The day of onset of symptoms was defined as day 0 of the illness. The last visit was performed at the time of discharge for patients who recovered entirely, or as a follow-up visit for patients still in the critical phase. A clinical and biological follow-up including abdominal/chest ultrasound recording was conducted at each visit. DENV infection of hospitalized patients was confirmed by NS1 antigen detection using our NS1-capture ELISA (*47–49*) and/or RT-qPCR and/or virus isolation on *Aedes albopictus* C6/36 cells on the plasma sample obtained at admission (*85*). Finally, the severity of the disease among confirmed dengue patients was assessed according to the WHO 2009 criteria using clinical and biological data recorded at admission (ADM), follow up visit (F-VIS) or discharge (DIS).

### Capture ELISA of NS1-HDL complexes

Microtitration plates were coated overnight with purified mouse anti-NS1 monoclonal antibody. Wells were saturated and washed before serial dilutions of human sera spiked with purified DENV-2 NS1 or DENV1- or DENV-2-infected human sera were added to wells for 2 h at room temperature. Wells were washed again and incubated for 1 h at 37°C with anti-ApoA-I goat polyclonal antibodies followed by a peroxidase-conjugated secondary antibody revealed with a 3,3”, 5,5”-tetramethylbenzidine solution. Negative controls were measured when the reaction was carried out in the absence of antigen. Absorbance values were corrected by substracting the mean value of the signals measured for the negative controls.

## ACKNOWLEDGMENTS

The authors gratefully acknowledge the staff of the Kampong Cham Referral Hospital, the patients and parents who participated in the study, and the Arbovirus Team in the Virology Unit at the Institut Pasteur du Cambodge who contributed to this study. We acknowledge the contribution of the ICAReB facility in setting up the recruitment of donors and the acquisition of blood samples, in particular Gloria Morizot, Bianca Liliana Perlaza, Sophie Chaouche, Linda Sangari, Céline Chapel, Philippe Esterre and Hélène Laude. We are most grateful to Christine Girard-Blanc and Evelyne Dufour for their contribution in producing and purifying the recombinant DENV NS1 protein. We thank M. Nilges and the Equipex CACSICE for providing the Falcon II direct detector and David Vreesler for his help in acquiring the first electron microscopy images of the bovine NS1-HDL complex. Finally, we thank Alexandre Pachot and Karine Kaiser for their support and helpful discussions.

## Ethics Statement

The study on dengue virus-infected patients was approved by the Cambodian National Ethics Committee for Human Research (approval #087NECHR/2011). All patient inclusion and blood sampling occurred after obtaining written informed consent from the patient’s parents or guardians.

Primary monocyte-derived macrophages were isolated from healthy donor blood obtained from the French blood bank (Etablissement Français du Sang) as part of a convention with the Institut Pasteur. In accordance with French law, written informed consent to use the cells for clinical research was obtained from each donor.

