## Supplemental Figure 1 for "The dengue virus NS1 protein conveys pro-inflammatory signals by docking onto human high-density lipoproteins"

### Supplementary Fig. 1

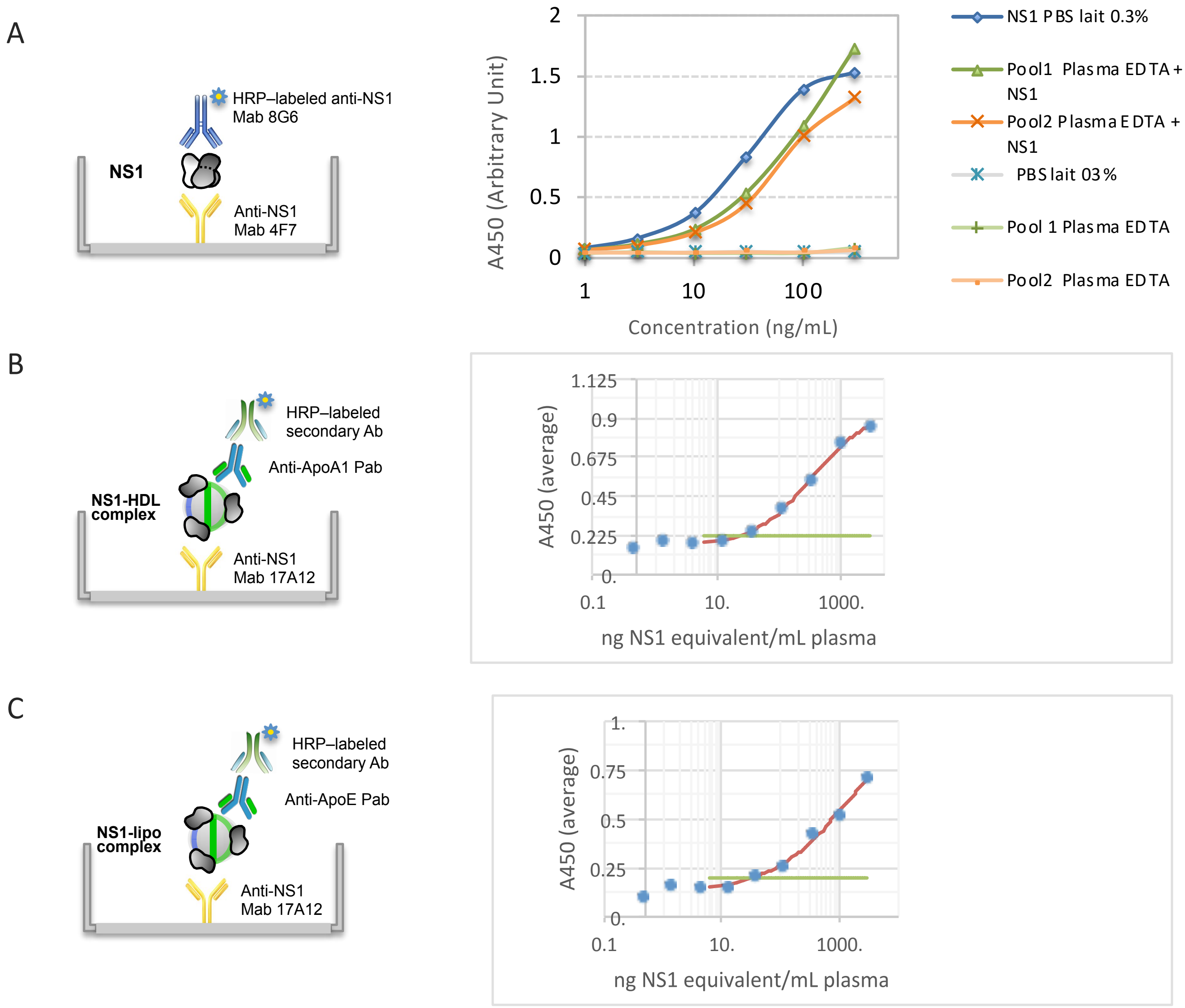


Supplementary Fig. 1: Standard calibration curves for the detection of the DENV NS1 protein alone (A), the NS1-ApoA-I complex (B) or the NS1-ApoE complex (C).

(A) Detection of the purified DENV NS1 has been described previously (Alcon-LePoder, 2006). The principle of the assay is presented in the left-hand side diagram. (B, C) NS1-ApoA-I or NS1-ApoE complexes were assembled *in vitro* by incubating a purified preparation of NS1 at a known concentration in normal plasma obtained from a human donor (ICAReB Facility, I. Pasteur). Capture of the NS1-ApoA-I or NS1-ApoE complexes was carried out using an anti-NS1 monoclonal antibody (MAb17A12). The detection of immobilized complexes was performed with an anti-ApoAI or anti-ApoE polyclonal antibody followed by a species-specific secondary antibody. The concentration of the NS1-ApoA-I or NS1-ApoE complexes is reported on the basis of 100% NS1 bound to HDL (B) or ApoE-positive lipoprotein particles (C), respectively. Detection limits of the NS1, NS1-ApoA-I or NS1-ApoE assays were set as twice the mean value of signals obtained with normal human plasma in the absence of NS1, which corresponded to 0.5, 17 and 15 ng of an equivalent NS1 concentration per milliliter, respectively.
